## Supplementary File 1 for "Boosting biodiversity monitoring using smartphone-driven, rapidly accumulating community-sourced data": S1_file.html

S1 File: Distributions of occurrence records along with environmental variables


Code 

- Show All Code
- Hide All Code

### S1 File: Distributions of occurrence records along with environmental variables

```
rm(list = ls())

pacman::p_load(
  tidyverse, data.table, 
  gridExtra, patchwork, cowplot, grid, ggbreak,
  conflicted,
  knitr, kableExtra, pander  # nice tables
  )

opts_chunk$set(
  prompt=TRUE, message=FALSE, comment="", warning = FALSE
  ) 

options(
  knitr.kable.NA = '',
  scipen=100           # do not show numbers using exponential
  )

conflicted::conflict_prefer_all(winner = 'dplyr', quiet = TRUE)

changeVarNames = function(varnames) {
  varnames_out = varnames %>% 
    str_replace_all(., "^x$", "longitude") %>% 
    str_replace_all(., "^y$", "latitude") %>% 
    str_replace_all(., "city", "urban") %>% 
    str_replace_all(., "plantation", "planted forest") %>% 
    str_replace_all(., "cliamte", "bioclim 15") %>% 
    str_replace_all(., "rice", "rice fild") %>% 
    str_replace_all(., "grass", "seminatural") %>%
    str_replace_all(., "hokkaido", "Hokkaido") %>% 
    str_replace_all(., '_', ' ')
}
```

```
> ## Load occurrence count data 
> df = fread('data/product/dataDistribution.csv', encoding = 'UTF-8') %>% 
+   select(-mesh_code_3rd)
> 
> ## Select columns with environmental variables
> cols = colnames(df) %>% str_subset('postCount', negate=T)
> 
> ## Draw occurrence records' histograms for each environmental variable 
> for(coln in 1:length(cols)) {
+   ## Select 1 environmental variable and post count data 
+   cols_need = c(cols[coln], 'postCount_biome', 'postCount_expert')
+   df_counts = df %>% 
+     select(one_of(cols_need))
+   colnames(df_counts) = c('envVar', 'postCount_biome', 'postCount_expert')
+   
+   ## Prepare histogram
+   df_counts = df_counts %>% 
+     ## Based on env condition, split dataset into detemined n of bins
+     mutate(
+       cut = cut_interval(envVar, n = 20) %>% 
+         as.character()
+     ) %>% 
+     group_by(cut) %>% 
+     summarise(across(starts_with('postCount_'), sum), .groups = 'drop') %>% 
+     mutate(
+       bin_start = cut %>% 
+         str_remove_all(., '\\[|\\]|\\(|,.*') %>% 
+         as.numeric(),
+       bin_end = cut %>% 
+         str_remove_all(., '\\[|\\]|\\)|.*,') %>% 
+         as.numeric(),
+       bin_mean = round((bin_start + bin_end)/2, 2)
+     )
+   
+   binwidth = (df_counts$bin_end[1] - df_counts$bin_start[1])*0.5
+   
+   ## Visualize histogram
+   p = df_counts %>% 
+     ggplot() +
+     geom_bar(
+       aes(x = bin_mean, y = postCount_biome), 
+       stat = 'identity', fill = '#006837', alpha = 0.3, width = binwidth*1.2
+     ) +
+     geom_bar(
+       aes(x = bin_mean, y = postCount_expert), 
+       stat = 'identity', fill = 'grey30', alpha = 0.3, width = binwidth #coral
+     ) +
+     labs(x = NULL, y = NULL, fill = 'legend') +
+     ggtitle(changeVarNames(cols[coln]))
+   
+   assign(str_c('p', coln), p)
+ }
> 
> ## Prepare shared x&y labels
> xlab = 'Standardised value'
> ylab = 'Occurrence records'
```

Distributions of occurrence records of *Biome* (green colour) and Traditional survey data
( grey colour) along with
environmental variables are shown.

### Data distibution against principal components of all variables used

```
> p_allPC = (
+   (p55+p56+p57+p58+p59+p60+p61+p62+p63+p64+p65+
+      plot_layout(guides = "collect", ncol = 3)
+    )
+   & labs(x = NULL, y = NULL)
+   & theme(
+     plot.title = element_text(hjust = 1, vjust = -8, size = 15),
+     axis.text  = element_text(size = 11)
+     )
+   )
> grid.arrange(
+   patchworkGrob(p_allPC), 
+   left = ylab, 
+   bottom = str_c(xlab, "\nFigure S1-1: Occurrence records distributions against PCs of all environmental variables. The PCs explaining >2% of total variance are shown"),
+   top = NULL
+   )
```

```
> fread('data/product/dataQuality/envVars_PCimportance.csv', encoding = 'UTF-8') %>% 
+   kable("html", digits = 1, caption = "S1-1 Table. cumulative importance of PCs") %>%
+   kable_styling("striped", position = "left") %>% 
+   scroll_box(height = "400px")
```

S1-1 Table. cumulative importance of PCs

| name | cumulative imoprtance (%) | proportion of variance (%) |
| --- | --- | --- |
| PC1 | 6.1 | 6.1 |
| PC2 | 11.0 | 4.9 |
| PC3 | 14.0 | 3.0 |
| PC4 | 16.7 | 2.7 |
| PC5 | 19.3 | 2.6 |
| PC6 | 21.7 | 2.3 |
| PC7 | 24.0 | 2.3 |
| PC8 | 26.2 | 2.2 |
| PC9 | 28.3 | 2.1 |
| PC10 | 30.3 | 2.1 |
| PC11 | 32.4 | 2.0 |
| PC12 | 34.4 | 2.0 |
| PC13 | 36.3 | 2.0 |
| PC14 | 38.3 | 2.0 |
| PC15 | 40.3 | 2.0 |
| PC16 | 42.2 | 1.9 |
| PC17 | 44.1 | 1.9 |
| PC18 | 46.1 | 1.9 |
| PC19 | 48.0 | 1.9 |
| PC20 | 49.9 | 1.9 |
| PC21 | 51.9 | 1.9 |
| PC22 | 53.8 | 1.9 |
| PC23 | 55.7 | 1.9 |
| PC24 | 57.7 | 1.9 |
| PC25 | 59.6 | 1.9 |
| PC26 | 61.5 | 1.9 |
| PC27 | 63.5 | 1.9 |
| PC28 | 65.4 | 1.9 |
| PC29 | 67.3 | 1.9 |
| PC30 | 69.2 | 1.9 |
| PC31 | 71.2 | 1.9 |
| PC32 | 73.1 | 1.9 |
| PC33 | 75.0 | 1.9 |
| PC34 | 76.9 | 1.9 |
| PC35 | 78.9 | 1.9 |
| PC36 | 80.8 | 1.9 |
| PC37 | 82.7 | 1.9 |
| PC38 | 84.6 | 1.9 |
| PC39 | 86.4 | 1.8 |
| PC40 | 88.2 | 1.8 |
| PC41 | 89.9 | 1.7 |
| PC42 | 91.4 | 1.5 |
| PC43 | 92.9 | 1.5 |
| PC44 | 94.3 | 1.4 |
| PC45 | 95.6 | 1.3 |
| PC46 | 96.6 | 1.0 |
| PC47 | 97.5 | 0.9 |
| PC48 | 98.3 | 0.8 |
| PC49 | 99.0 | 0.7 |
| PC50 | 99.4 | 0.4 |
| PC51 | 99.8 | 0.4 |
| PC52 | 100.0 | 0.2 |

```
> fread('data/product/dataQuality/envVars_PCrotation.csv', encoding = 'UTF-8') %>% 
+   mutate_at('Environmental variables', changeVarNames) %>% 
+   kable("html", digits = 2, caption = "S1-2 Table. variable contributions to PCs") %>%
+   kable_styling("striped", position = "left") %>% 
+   scroll_box(height = "400px")
```

S1-2 Table. variable contributions to PCs

| Environmental variables | PC1 | PC2 | PC3 | PC4 | PC5 | PC6 | PC7 | PC8 | PC9 | PC10 | PC11 | PC12 | PC13 | PC14 | PC15 | PC16 | PC17 | PC18 | PC19 | PC20 | PC21 | PC22 | PC23 | PC24 | PC25 | PC26 | PC27 | PC28 | PC29 | PC30 | PC31 | PC32 | PC33 | PC34 | PC35 | PC36 | PC37 | PC38 | PC39 | PC40 | PC41 | PC42 | PC43 | PC44 | PC45 | PC46 | PC47 | PC48 | PC49 | PC50 | PC51 | PC52 |
| --- | --- | --- | --- | --- | --- | --- | --- | --- | --- | --- | --- | --- | --- | --- | --- | --- | --- | --- | --- | --- | --- | --- | --- | --- | --- | --- | --- | --- | --- | --- | --- | --- | --- | --- | --- | --- | --- | --- | --- | --- | --- | --- | --- | --- | --- | --- | --- | --- | --- | --- | --- | --- |
| PC1 land use | 0.45 | -0.07 | 0.02 | -0.02 | 0.17 | -0.02 | 0.10 | -0.01 | 0.12 | -0.10 | 0.04 | -0.12 | -0.04 | -0.03 | 0.04 | 0.01 | 0.01 | 0.00 | 0.00 | 0.00 | 0.00 | 0.00 | 0.01 | 0.01 | 0.01 | 0.00 | 0.01 | 0.00 | 0.00 | 0.00 | 0.00 | 0.01 | 0.01 | 0.02 | 0.01 | 0.00 | 0.01 | -0.02 | -0.01 | 0.01 | -0.15 | 0.08 | -0.17 | 0.09 | -0.03 | 0.19 | -0.04 | 0.22 | -0.55 | 0.29 | -0.39 | 0.13 |
| PC2 land use | -0.01 | -0.05 | -0.34 | 0.17 | 0.23 | 0.06 | 0.23 | 0.39 | 0.25 | -0.07 | -0.13 | 0.10 | 0.00 | 0.03 | 0.01 | 0.01 | 0.02 | 0.00 | -0.01 | 0.00 | 0.01 | 0.00 | -0.01 | 0.00 | 0.00 | 0.00 | 0.00 | 0.00 | 0.00 | -0.01 | 0.01 | 0.02 | 0.01 | -0.03 | 0.00 | 0.02 | 0.04 | -0.01 | -0.03 | -0.05 | -0.15 | -0.05 | -0.45 | -0.44 | 0.06 | -0.18 | 0.06 | 0.11 | 0.12 | -0.04 | 0.04 | -0.08 |
| PC3 land use | -0.06 | -0.23 | 0.03 | 0.34 | 0.12 | -0.30 | 0.14 | -0.02 | -0.24 | -0.29 | 0.09 | 0.00 | -0.02 | 0.03 | -0.06 | -0.03 | -0.01 | 0.00 | 0.00 | 0.00 | -0.01 | 0.00 | 0.00 | 0.00 | -0.01 | 0.00 | 0.00 | 0.01 | 0.00 | 0.00 | 0.00 | 0.00 | 0.00 | 0.01 | 0.01 | 0.01 | 0.03 | -0.01 | -0.01 | -0.47 | 0.15 | -0.12 | -0.01 | 0.19 | -0.04 | 0.19 | 0.01 | 0.33 | 0.29 | 0.11 | -0.05 | -0.03 |
| PC4 land use | 0.05 | -0.08 | -0.13 | 0.22 | -0.22 | 0.08 | -0.23 | 0.15 | -0.29 | 0.25 | -0.42 | -0.30 | -0.06 | -0.09 | -0.03 | 0.04 | -0.01 | -0.02 | -0.01 | -0.02 | 0.01 | 0.00 | 0.00 | 0.00 | 0.02 | -0.02 | -0.01 | -0.01 | -0.01 | 0.01 | -0.01 | -0.06 | -0.01 | -0.07 | -0.05 | -0.06 | -0.24 | 0.07 | -0.01 | 0.10 | -0.33 | 0.00 | 0.09 | 0.06 | 0.00 | -0.07 | -0.02 | 0.38 | 0.07 | 0.03 | -0.01 | 0.02 |
| PC5 land use | 0.00 | 0.03 | 0.11 | -0.33 | 0.28 | -0.16 | 0.12 | 0.24 | -0.47 | 0.08 | -0.32 | -0.11 | -0.02 | -0.04 | -0.06 | 0.02 | -0.02 | -0.02 | -0.02 | -0.01 | 0.00 | 0.00 | 0.00 | 0.00 | 0.01 | -0.02 | -0.01 | -0.01 | -0.01 | 0.01 | -0.01 | -0.05 | -0.01 | -0.08 | -0.05 | -0.04 | -0.19 | 0.03 | 0.00 | -0.26 | 0.06 | -0.07 | -0.15 | -0.05 | 0.01 | 0.06 | 0.02 | -0.42 | -0.13 | 0.01 | -0.04 | -0.01 |
| PC6 land use | 0.01 | 0.07 | -0.09 | 0.03 | -0.10 | 0.15 | 0.55 | -0.24 | -0.24 | 0.24 | -0.04 | 0.01 | 0.01 | -0.03 | 0.02 | 0.02 | -0.01 | -0.01 | -0.01 | 0.00 | 0.00 | 0.00 | 0.00 | 0.00 | 0.00 | -0.01 | 0.00 | 0.00 | 0.00 | 0.00 | 0.00 | -0.03 | -0.01 | -0.03 | -0.02 | -0.02 | -0.08 | 0.00 | -0.01 | 0.19 | 0.39 | 0.34 | -0.31 | 0.15 | -0.01 | -0.05 | 0.00 | 0.20 | 0.01 | -0.07 | 0.04 | 0.01 |
| PC1 climate | 0.01 | -0.55 | 0.05 | -0.11 | -0.04 | 0.05 | 0.00 | -0.14 | 0.02 | -0.10 | -0.12 | 0.04 | 0.01 | 0.00 | 0.01 | 0.00 | 0.00 | 0.00 | 0.00 | 0.00 | 0.00 | 0.00 | 0.00 | 0.00 | 0.00 | 0.00 | 0.00 | 0.00 | 0.00 | 0.00 | 0.00 | 0.00 | 0.00 | 0.00 | 0.00 | 0.00 | 0.00 | 0.00 | -0.01 | -0.03 | 0.00 | 0.03 | 0.01 | 0.00 | 0.05 | -0.17 | -0.30 | 0.03 | -0.26 | -0.10 | 0.17 | -0.63 |
| PC2 climate | 0.34 | -0.07 | -0.34 | -0.12 | -0.21 | -0.07 | -0.06 | 0.11 | -0.12 | 0.01 | 0.11 | 0.06 | 0.06 | -0.01 | 0.04 | 0.01 | 0.01 | 0.00 | 0.00 | 0.00 | 0.00 | 0.00 | 0.00 | 0.00 | 0.00 | 0.00 | 0.01 | 0.00 | 0.00 | 0.00 | 0.00 | 0.00 | 0.00 | 0.00 | 0.00 | 0.00 | 0.01 | 0.01 | 0.02 | 0.07 | 0.12 | -0.08 | 0.07 | -0.11 | -0.05 | 0.46 | 0.50 | 0.03 | -0.04 | -0.22 | 0.12 | -0.26 |
| PC3 climate | 0.26 | -0.01 | 0.46 | 0.31 | 0.11 | 0.03 | -0.05 | 0.05 | -0.06 | 0.03 | 0.00 | 0.09 | 0.04 | -0.01 | 0.02 | 0.01 | 0.00 | 0.00 | 0.00 | 0.00 | 0.00 | 0.00 | 0.00 | 0.00 | 0.00 | 0.00 | 0.00 | 0.00 | 0.00 | 0.00 | 0.00 | 0.00 | -0.01 | 0.00 | 0.00 | 0.00 | 0.00 | -0.02 | 0.00 | -0.02 | 0.07 | 0.00 | 0.03 | 0.04 | 0.02 | -0.50 | 0.55 | -0.02 | -0.10 | -0.13 | -0.04 | -0.10 |
| elevation diff | 0.51 | 0.01 | -0.02 | 0.03 | 0.03 | 0.01 | 0.00 | -0.05 | 0.04 | 0.01 | -0.04 | 0.02 | -0.03 | 0.03 | -0.06 | -0.02 | 0.00 | 0.01 | 0.00 | 0.00 | -0.01 | 0.00 | 0.01 | 0.00 | -0.01 | 0.00 | 0.00 | 0.01 | 0.00 | 0.00 | 0.00 | 0.00 | 0.00 | 0.00 | 0.00 | 0.00 | 0.01 | 0.00 | -0.01 | -0.07 | -0.07 | 0.05 | -0.05 | 0.12 | -0.01 | -0.05 | -0.09 | -0.14 | 0.15 | 0.38 | 0.69 | 0.13 |
| slope max | 0.47 | 0.01 | 0.00 | 0.01 | 0.04 | 0.02 | -0.01 | -0.05 | 0.08 | 0.01 | -0.02 | -0.05 | -0.05 | 0.01 | -0.04 | -0.01 | -0.01 | 0.00 | 0.01 | 0.00 | 0.00 | 0.00 | -0.01 | -0.01 | 0.00 | 0.00 | -0.01 | 0.00 | 0.00 | -0.01 | 0.00 | 0.00 | 0.01 | 0.01 | 0.01 | 0.01 | 0.01 | -0.01 | -0.01 | -0.05 | -0.16 | 0.12 | -0.10 | 0.20 | -0.03 | 0.01 | -0.27 | -0.21 | 0.44 | -0.48 | -0.33 | -0.07 |
| PC1 vegetation | 0.10 | -0.25 | -0.28 | 0.10 | -0.22 | 0.11 | 0.15 | -0.14 | 0.01 | 0.21 | -0.03 | 0.08 | -0.11 | 0.15 | -0.16 | -0.09 | -0.02 | 0.11 | 0.05 | -0.01 | -0.04 | 0.04 | -0.03 | -0.02 | -0.03 | -0.01 | -0.12 | 0.03 | 0.04 | -0.26 | 0.22 | -0.23 | -0.07 | -0.03 | 0.02 | -0.02 | 0.02 | -0.02 | 0.08 | -0.17 | 0.12 | -0.18 | 0.21 | -0.13 | 0.04 | -0.18 | 0.05 | -0.26 | 0.01 | 0.23 | -0.29 | 0.11 |
| PC2 vegetation | 0.08 | -0.26 | -0.20 | -0.23 | -0.14 | -0.07 | -0.19 | 0.11 | -0.16 | -0.14 | 0.04 | 0.17 | 0.08 | -0.03 | 0.03 | -0.04 | 0.01 | -0.09 | 0.01 | 0.03 | 0.01 | -0.03 | 0.03 | -0.07 | 0.09 | -0.02 | -0.03 | 0.08 | -0.01 | 0.33 | -0.33 | 0.22 | 0.09 | -0.01 | 0.06 | -0.02 | 0.01 | 0.07 | -0.16 | 0.12 | 0.19 | 0.13 | -0.03 | 0.01 | 0.06 | -0.30 | 0.03 | -0.03 | 0.18 | 0.25 | -0.23 | 0.15 |
| PC3 vegetation | 0.24 | 0.18 | 0.05 | 0.00 | -0.13 | -0.02 | 0.03 | 0.02 | -0.22 | -0.35 | -0.11 | 0.01 | 0.31 | -0.10 | 0.06 | 0.02 | -0.04 | -0.07 | 0.09 | 0.16 | -0.06 | -0.08 | -0.02 | -0.04 | -0.06 | 0.02 | -0.13 | 0.08 | 0.02 | -0.12 | 0.20 | 0.01 | 0.01 | 0.02 | 0.01 | 0.01 | 0.02 | -0.08 | 0.06 | 0.04 | 0.19 | -0.11 | 0.18 | -0.33 | 0.20 | -0.11 | -0.29 | 0.17 | -0.09 | -0.19 | 0.05 | 0.20 |
| PC4 vegetation | 0.05 | 0.31 | -0.31 | -0.10 | -0.03 | -0.10 | 0.00 | 0.02 | -0.02 | 0.18 | 0.19 | 0.11 | -0.02 | -0.03 | -0.12 | -0.04 | 0.14 | -0.10 | -0.13 | -0.17 | -0.13 | 0.03 | 0.00 | -0.03 | 0.06 | 0.02 | -0.10 | -0.02 | -0.02 | 0.04 | -0.20 | -0.01 | -0.01 | 0.00 | 0.09 | 0.02 | 0.02 | 0.06 | -0.02 | -0.35 | -0.02 | -0.22 | 0.05 | 0.20 | -0.11 | -0.32 | -0.13 | 0.18 | -0.30 | -0.24 | 0.08 | 0.00 |
| PC5 vegetation | -0.09 | 0.02 | -0.23 | 0.34 | 0.03 | -0.14 | -0.12 | 0.00 | -0.08 | -0.14 | -0.08 | -0.12 | 0.02 | 0.08 | -0.13 | -0.01 | 0.20 | -0.01 | 0.20 | 0.32 | -0.29 | -0.18 | 0.16 | -0.02 | -0.14 | 0.13 | -0.02 | -0.06 | -0.01 | 0.04 | -0.04 | 0.01 | -0.06 | 0.12 | 0.05 | -0.03 | 0.06 | -0.06 | 0.13 | 0.26 | 0.06 | -0.06 | -0.17 | 0.21 | -0.22 | -0.04 | -0.03 | -0.28 | -0.17 | -0.03 | 0.02 | -0.03 |
| PC6 vegetation | 0.13 | 0.00 | 0.09 | 0.10 | 0.29 | -0.04 | 0.05 | 0.22 | 0.06 | 0.37 | -0.04 | 0.10 | -0.03 | 0.04 | -0.09 | -0.02 | -0.07 | 0.23 | 0.03 | 0.08 | 0.00 | 0.12 | -0.04 | 0.07 | -0.07 | 0.07 | -0.04 | -0.03 | -0.07 | 0.37 | -0.06 | 0.05 | -0.18 | 0.01 | -0.01 | 0.05 | 0.08 | -0.01 | 0.03 | 0.22 | 0.30 | -0.23 | 0.27 | -0.08 | -0.05 | 0.10 | -0.24 | 0.13 | 0.04 | 0.04 | -0.04 | -0.09 |
| PC7 vegetation | -0.02 | 0.02 | 0.06 | 0.46 | -0.23 | 0.09 | -0.02 | 0.27 | -0.04 | 0.09 | 0.02 | 0.14 | 0.09 | 0.03 | -0.03 | -0.03 | 0.08 | -0.17 | -0.13 | -0.22 | 0.13 | 0.05 | 0.12 | -0.12 | 0.09 | -0.31 | -0.05 | 0.08 | -0.07 | 0.03 | 0.00 | 0.21 | -0.05 | 0.06 | 0.07 | 0.04 | 0.00 | -0.01 | -0.09 | -0.09 | 0.08 | 0.18 | 0.00 | 0.00 | 0.13 | 0.22 | -0.19 | -0.28 | -0.20 | 0.00 | -0.01 | -0.02 |
| PC8 vegetation | 0.00 | 0.06 | 0.03 | 0.01 | -0.11 | 0.10 | 0.08 | -0.42 | 0.15 | -0.18 | -0.45 | 0.09 | -0.04 | 0.13 | -0.15 | -0.01 | -0.08 | 0.11 | -0.08 | -0.04 | -0.03 | 0.01 | 0.07 | -0.05 | -0.13 | -0.21 | -0.06 | 0.02 | -0.09 | 0.41 | -0.18 | 0.14 | -0.04 | 0.03 | -0.07 | -0.09 | -0.01 | -0.01 | 0.07 | -0.15 | -0.07 | -0.16 | -0.04 | -0.10 | 0.01 | 0.11 | 0.15 | -0.01 | -0.07 | -0.12 | 0.00 | 0.13 |
| PC9 vegetation | -0.06 | -0.05 | -0.12 | 0.02 | 0.07 | 0.09 | 0.22 | 0.05 | 0.23 | -0.07 | -0.08 | -0.21 | 0.50 | -0.39 | -0.21 | 0.05 | -0.12 | 0.04 | -0.01 | 0.00 | -0.03 | 0.02 | 0.00 | -0.02 | -0.01 | -0.02 | -0.03 | 0.01 | -0.02 | 0.02 | -0.02 | -0.07 | -0.03 | -0.04 | -0.04 | 0.02 | -0.06 | -0.12 | -0.42 | -0.02 | -0.02 | -0.01 | 0.23 | 0.19 | -0.13 | -0.01 | 0.11 | -0.12 | 0.02 | 0.03 | 0.00 | -0.02 |
| PC10 vegetation | 0.03 | -0.08 | 0.13 | -0.05 | 0.05 | -0.08 | -0.10 | -0.16 | 0.04 | 0.18 | 0.14 | 0.01 | 0.30 | -0.10 | -0.58 | 0.17 | 0.25 | -0.29 | -0.11 | -0.04 | 0.10 | -0.04 | 0.07 | 0.04 | 0.05 | -0.01 | 0.03 | -0.03 | 0.02 | 0.01 | 0.00 | -0.04 | 0.05 | -0.03 | -0.01 | -0.09 | -0.09 | 0.00 | 0.31 | 0.09 | -0.01 | -0.09 | -0.17 | -0.17 | 0.07 | 0.05 | -0.01 | 0.07 | 0.10 | 0.08 | -0.05 | -0.03 |
| PC11 vegetation | -0.06 | 0.04 | 0.01 | -0.15 | -0.07 | 0.16 | 0.10 | 0.18 | -0.09 | 0.05 | -0.04 | 0.23 | 0.18 | 0.14 | -0.28 | -0.03 | -0.04 | 0.16 | 0.46 | 0.26 | 0.10 | 0.04 | -0.04 | 0.13 | 0.12 | 0.02 | 0.07 | 0.05 | -0.02 | 0.00 | 0.01 | 0.18 | -0.01 | 0.10 | 0.10 | 0.14 | 0.11 | -0.05 | 0.20 | -0.21 | -0.26 | 0.25 | 0.09 | 0.14 | 0.02 | 0.01 | 0.06 | 0.09 | -0.05 | -0.01 | 0.00 | -0.01 |
| PC12 vegetation | -0.05 | -0.02 | 0.07 | -0.18 | 0.10 | 0.05 | -0.03 | 0.13 | -0.01 | -0.08 | 0.00 | 0.12 | -0.07 | 0.10 | -0.22 | -0.13 | -0.04 | -0.05 | -0.14 | -0.15 | -0.20 | -0.06 | 0.10 | -0.20 | -0.32 | 0.06 | -0.56 | 0.06 | -0.08 | -0.12 | 0.20 | 0.15 | -0.21 | -0.15 | 0.04 | 0.00 | 0.01 | 0.06 | -0.02 | 0.11 | -0.07 | 0.21 | 0.03 | 0.09 | -0.05 | 0.04 | 0.09 | 0.14 | 0.06 | 0.02 | 0.01 | -0.04 |
| PC13 vegetation | 0.02 | 0.04 | 0.04 | -0.07 | 0.04 | 0.03 | -0.15 | -0.01 | 0.14 | 0.06 | -0.27 | 0.13 | -0.08 | 0.07 | -0.06 | -0.06 | 0.04 | -0.07 | -0.04 | 0.12 | 0.30 | -0.34 | 0.43 | -0.06 | 0.27 | 0.22 | 0.11 | 0.19 | 0.10 | -0.18 | -0.01 | -0.08 | -0.22 | -0.05 | -0.03 | -0.02 | 0.12 | 0.00 | -0.25 | -0.08 | 0.20 | -0.03 | -0.02 | 0.05 | -0.05 | 0.06 | 0.01 | 0.12 | -0.01 | -0.04 | 0.00 | 0.03 |
| PC14 vegetation | 0.03 | 0.01 | 0.00 | -0.01 | 0.12 | 0.13 | 0.24 | 0.13 | -0.11 | -0.26 | -0.04 | 0.07 | -0.16 | 0.26 | -0.26 | -0.13 | 0.06 | -0.13 | -0.15 | -0.24 | -0.21 | -0.05 | 0.12 | 0.14 | 0.17 | 0.04 | 0.31 | -0.07 | 0.10 | 0.07 | 0.05 | -0.12 | 0.24 | 0.10 | 0.09 | -0.12 | -0.05 | 0.02 | -0.09 | 0.22 | -0.07 | 0.08 | 0.34 | -0.04 | -0.09 | 0.00 | 0.00 | 0.02 | 0.00 | -0.05 | 0.00 | 0.02 |
| PC15 vegetation | -0.02 | 0.01 | -0.05 | 0.02 | -0.03 | -0.14 | 0.01 | -0.04 | -0.01 | 0.03 | 0.09 | 0.26 | 0.05 | -0.04 | 0.03 | 0.02 | -0.06 | 0.02 | -0.05 | 0.06 | -0.11 | 0.01 | 0.12 | 0.02 | -0.16 | -0.05 | 0.23 | 0.44 | 0.29 | 0.24 | 0.36 | -0.04 | -0.02 | -0.09 | -0.17 | 0.29 | -0.27 | 0.20 | -0.03 | 0.05 | -0.14 | -0.08 | -0.07 | 0.12 | 0.03 | -0.03 | -0.02 | -0.02 | -0.03 | 0.00 | -0.02 | 0.00 |
| PC16 vegetation | 0.00 | 0.03 | 0.00 | 0.00 | 0.03 | 0.19 | -0.17 | 0.09 | 0.15 | 0.04 | -0.12 | -0.05 | -0.04 | 0.15 | -0.05 | -0.18 | 0.12 | -0.21 | -0.17 | 0.31 | 0.04 | -0.08 | -0.29 | 0.08 | -0.18 | -0.07 | -0.05 | 0.04 | 0.01 | 0.06 | 0.01 | -0.10 | 0.46 | -0.25 | -0.08 | 0.25 | -0.05 | -0.14 | -0.01 | -0.17 | 0.24 | 0.12 | 0.03 | 0.04 | -0.13 | 0.06 | 0.00 | 0.04 | -0.03 | -0.01 | 0.00 | 0.00 |
| PC17 vegetation | 0.01 | 0.02 | 0.05 | 0.06 | 0.03 | -0.01 | 0.06 | -0.12 | -0.04 | 0.15 | 0.17 | -0.09 | 0.01 | 0.15 | -0.07 | -0.12 | 0.01 | 0.19 | 0.16 | -0.01 | -0.07 | -0.19 | -0.05 | -0.43 | 0.25 | 0.14 | -0.12 | -0.04 | 0.20 | 0.01 | 0.03 | 0.37 | 0.31 | -0.03 | -0.26 | -0.07 | -0.22 | -0.15 | -0.18 | 0.00 | -0.08 | -0.12 | -0.01 | -0.12 | 0.01 | 0.04 | -0.02 | -0.01 | -0.02 | -0.01 | 0.02 | -0.05 |
| PC18 vegetation | -0.01 | -0.01 | -0.06 | 0.00 | 0.05 | -0.08 | -0.02 | -0.01 | 0.09 | -0.15 | -0.24 | 0.12 | 0.05 | 0.13 | 0.03 | 0.04 | 0.10 | -0.14 | -0.01 | -0.14 | 0.21 | 0.42 | -0.15 | -0.11 | 0.17 | 0.29 | -0.16 | -0.04 | -0.02 | -0.10 | 0.00 | -0.01 | -0.01 | 0.38 | -0.20 | 0.31 | -0.24 | -0.02 | 0.08 | 0.13 | 0.09 | -0.05 | 0.01 | 0.14 | -0.06 | -0.01 | 0.04 | -0.01 | 0.00 | 0.00 | -0.01 | 0.05 |
| PC19 vegetation | -0.02 | 0.02 | 0.10 | 0.02 | -0.05 | -0.04 | -0.13 | -0.08 | -0.07 | 0.03 | -0.03 | 0.12 | 0.06 | 0.06 | -0.09 | -0.12 | -0.01 | 0.06 | -0.20 | 0.35 | -0.36 | 0.55 | -0.01 | 0.04 | 0.32 | 0.00 | -0.02 | 0.02 | 0.01 | -0.10 | 0.01 | -0.06 | -0.03 | -0.15 | 0.04 | -0.14 | 0.12 | -0.01 | -0.28 | 0.03 | -0.05 | 0.00 | -0.18 | -0.05 | 0.06 | 0.09 | -0.01 | 0.03 | -0.01 | -0.02 | 0.00 | 0.01 |
| PC20 vegetation | 0.01 | -0.01 | -0.03 | -0.04 | 0.02 | 0.07 | -0.03 | 0.12 | 0.09 | -0.03 | 0.03 | 0.15 | -0.08 | -0.20 | -0.01 | 0.11 | -0.19 | 0.05 | -0.14 | 0.26 | -0.16 | -0.13 | 0.21 | -0.03 | 0.19 | -0.30 | -0.11 | -0.42 | 0.07 | -0.06 | 0.04 | -0.03 | 0.05 | 0.22 | -0.38 | 0.02 | -0.09 | 0.28 | 0.18 | -0.08 | 0.09 | 0.08 | 0.06 | 0.05 | -0.04 | -0.02 | -0.02 | 0.03 | 0.02 | 0.02 | -0.02 | 0.00 |
| PC21 vegetation | -0.01 | 0.00 | 0.04 | 0.04 | 0.06 | -0.07 | 0.05 | 0.00 | 0.12 | 0.03 | -0.05 | 0.27 | 0.02 | -0.04 | 0.04 | 0.07 | -0.16 | 0.04 | 0.01 | 0.17 | 0.02 | -0.08 | -0.05 | -0.39 | 0.02 | 0.00 | 0.03 | 0.04 | -0.29 | -0.10 | -0.08 | -0.19 | 0.23 | -0.01 | 0.53 | -0.10 | -0.36 | 0.18 | 0.05 | 0.08 | -0.02 | -0.08 | 0.02 | 0.08 | 0.02 | 0.08 | -0.01 | 0.01 | 0.00 | 0.00 | 0.00 | -0.01 |
| PC22 vegetation | -0.01 | 0.00 | 0.03 | 0.00 | 0.02 | 0.00 | 0.00 | -0.07 | 0.05 | -0.02 | -0.07 | -0.14 | -0.12 | -0.09 | -0.03 | 0.02 | -0.06 | -0.09 | 0.12 | -0.01 | 0.03 | 0.02 | -0.01 | 0.06 | 0.19 | -0.13 | -0.28 | -0.18 | 0.60 | 0.15 | 0.07 | 0.01 | -0.05 | -0.01 | 0.53 | 0.24 | -0.02 | 0.04 | -0.02 | 0.01 | 0.05 | -0.04 | -0.04 | -0.04 | 0.00 | 0.03 | 0.05 | -0.01 | 0.01 | -0.02 | 0.03 | -0.01 |
| PC23 vegetation | -0.01 | 0.00 | 0.02 | 0.09 | -0.06 | -0.16 | -0.11 | -0.12 | -0.15 | 0.06 | -0.08 | 0.24 | 0.10 | 0.02 | -0.12 | -0.01 | -0.27 | 0.16 | -0.03 | -0.24 | 0.00 | -0.28 | -0.10 | 0.36 | 0.05 | 0.08 | -0.10 | -0.22 | -0.20 | -0.10 | 0.02 | -0.02 | 0.22 | -0.04 | 0.00 | 0.34 | 0.14 | 0.06 | -0.24 | 0.15 | -0.08 | -0.11 | -0.19 | 0.00 | 0.04 | 0.04 | -0.04 | -0.03 | -0.05 | 0.00 | -0.03 | 0.03 |
| PC24 vegetation | 0.00 | 0.02 | 0.01 | -0.01 | -0.04 | -0.05 | -0.20 | 0.12 | 0.21 | 0.09 | -0.19 | 0.16 | 0.08 | -0.01 | -0.01 | 0.03 | -0.18 | 0.03 | 0.14 | -0.22 | -0.19 | 0.01 | -0.17 | 0.23 | -0.19 | 0.07 | 0.08 | 0.06 | 0.31 | -0.17 | 0.03 | 0.15 | 0.11 | 0.18 | 0.02 | -0.48 | -0.12 | -0.03 | 0.01 | -0.10 | 0.25 | 0.04 | -0.12 | 0.13 | 0.05 | 0.05 | -0.03 | 0.05 | -0.03 | 0.01 | -0.02 | 0.01 |
| PC25 vegetation | 0.02 | -0.01 | 0.02 | -0.07 | -0.04 | 0.16 | -0.13 | -0.01 | -0.09 | 0.04 | 0.05 | -0.14 | 0.00 | -0.06 | -0.08 | 0.12 | 0.19 | 0.44 | -0.35 | 0.05 | -0.03 | 0.02 | 0.16 | -0.01 | -0.22 | 0.13 | 0.12 | 0.01 | -0.01 | -0.04 | 0.08 | 0.14 | 0.08 | 0.45 | 0.25 | 0.18 | -0.08 | -0.04 | -0.13 | -0.16 | 0.07 | 0.09 | -0.07 | -0.16 | -0.04 | -0.03 | 0.00 | 0.04 | 0.03 | 0.02 | -0.02 | 0.01 |
| PC26 vegetation | 0.01 | -0.01 | -0.06 | 0.02 | 0.05 | 0.06 | 0.03 | -0.09 | 0.08 | -0.06 | 0.12 | -0.03 | -0.02 | 0.05 | 0.13 | -0.08 | 0.07 | -0.06 | -0.01 | 0.12 | 0.06 | -0.03 | 0.11 | 0.51 | 0.16 | 0.01 | -0.11 | -0.05 | -0.27 | -0.01 | 0.16 | 0.36 | -0.16 | -0.13 | 0.10 | -0.10 | -0.53 | -0.02 | -0.03 | -0.05 | 0.03 | -0.07 | 0.05 | -0.04 | -0.09 | -0.04 | 0.01 | -0.03 | 0.01 | 0.00 | 0.00 | 0.00 |
| PC27 vegetation | -0.02 | -0.01 | -0.03 | -0.02 | 0.05 | -0.18 | 0.23 | -0.01 | -0.11 | 0.10 | -0.02 | 0.14 | 0.03 | 0.14 | 0.11 | 0.11 | 0.14 | -0.06 | -0.25 | 0.30 | 0.35 | -0.05 | -0.05 | 0.15 | -0.17 | -0.18 | -0.24 | 0.04 | 0.19 | -0.07 | 0.00 | 0.07 | 0.15 | 0.23 | -0.01 | -0.27 | 0.15 | -0.05 | -0.17 | 0.16 | -0.25 | -0.15 | 0.11 | 0.14 | 0.05 | -0.03 | 0.01 | -0.01 | 0.01 | 0.01 | 0.00 | -0.02 |
| PC28 vegetation | 0.03 | -0.01 | 0.03 | 0.07 | 0.08 | 0.07 | 0.06 | -0.05 | -0.07 | -0.06 | 0.05 | 0.08 | 0.02 | 0.04 | -0.05 | -0.05 | -0.04 | 0.08 | -0.03 | 0.04 | -0.04 | 0.04 | 0.02 | 0.08 | -0.20 | -0.10 | 0.08 | 0.09 | 0.26 | -0.43 | -0.64 | 0.10 | -0.15 | -0.14 | -0.06 | 0.21 | -0.21 | 0.06 | 0.03 | 0.10 | -0.05 | -0.03 | 0.09 | -0.12 | -0.07 | 0.05 | -0.05 | -0.01 | 0.01 | 0.00 | -0.01 | 0.01 |
| PC29 vegetation | -0.01 | -0.01 | 0.08 | -0.08 | -0.14 | -0.26 | 0.14 | 0.06 | 0.14 | 0.10 | -0.05 | -0.16 | 0.02 | -0.15 | -0.02 | -0.75 | -0.02 | -0.04 | -0.07 | 0.06 | -0.03 | -0.04 | 0.09 | 0.03 | -0.05 | -0.02 | 0.08 | -0.04 | -0.04 | -0.03 | -0.05 | 0.02 | -0.01 | 0.22 | 0.04 | 0.09 | -0.03 | -0.03 | 0.12 | 0.05 | -0.03 | -0.06 | -0.04 | 0.06 | 0.33 | -0.02 | 0.05 | 0.02 | 0.03 | 0.00 | 0.01 | -0.03 |
| PC30 vegetation | 0.03 | -0.04 | 0.09 | -0.05 | 0.04 | 0.08 | 0.07 | -0.12 | -0.01 | 0.25 | -0.03 | 0.02 | -0.10 | 0.00 | 0.10 | 0.14 | -0.06 | -0.57 | 0.25 | 0.09 | -0.32 | 0.00 | 0.09 | -0.02 | -0.18 | 0.00 | 0.08 | -0.03 | -0.15 | -0.05 | -0.01 | 0.15 | 0.00 | 0.30 | 0.02 | 0.17 | 0.07 | 0.00 | -0.27 | -0.10 | -0.06 | -0.11 | 0.05 | -0.14 | 0.00 | 0.07 | 0.03 | 0.04 | 0.08 | 0.05 | -0.03 | -0.02 |
| PC31 vegetation | -0.01 | -0.04 | -0.03 | 0.08 | 0.23 | 0.28 | -0.09 | -0.19 | -0.20 | 0.06 | -0.05 | 0.28 | 0.14 | -0.39 | 0.21 | -0.40 | 0.17 | -0.01 | 0.07 | -0.10 | 0.06 | 0.04 | 0.02 | 0.00 | 0.01 | 0.05 | -0.08 | 0.04 | 0.06 | 0.02 | 0.05 | -0.01 | 0.12 | 0.07 | 0.02 | -0.12 | 0.12 | 0.12 | 0.11 | -0.03 | -0.11 | -0.02 | -0.01 | -0.11 | -0.37 | 0.09 | -0.03 | -0.02 | 0.04 | 0.04 | -0.02 | 0.01 |
| PC32 vegetation | 0.01 | -0.02 | -0.06 | -0.07 | 0.00 | -0.04 | -0.02 | -0.02 | -0.02 | 0.05 | -0.12 | 0.19 | 0.25 | 0.15 | 0.18 | -0.02 | 0.06 | -0.04 | -0.10 | -0.03 | 0.05 | 0.08 | 0.13 | -0.20 | -0.21 | 0.01 | 0.33 | -0.58 | 0.11 | -0.03 | 0.16 | 0.16 | -0.11 | -0.28 | 0.09 | 0.06 | 0.00 | -0.22 | 0.03 | -0.02 | -0.04 | -0.04 | 0.02 | 0.12 | -0.02 | -0.04 | -0.01 | 0.04 | 0.02 | 0.04 | -0.05 | 0.05 |
| PC33 vegetation | -0.02 | -0.07 | 0.05 | 0.03 | -0.05 | 0.06 | 0.14 | 0.02 | 0.03 | 0.10 | -0.10 | 0.07 | 0.19 | 0.02 | 0.25 | 0.12 | 0.04 | -0.05 | -0.38 | 0.03 | -0.34 | -0.33 | -0.26 | 0.04 | 0.26 | 0.30 | -0.05 | 0.11 | 0.08 | 0.08 | -0.11 | 0.03 | -0.19 | 0.03 | 0.05 | 0.06 | 0.05 | -0.11 | 0.26 | -0.09 | -0.13 | 0.05 | 0.08 | 0.04 | 0.08 | 0.09 | 0.03 | -0.07 | 0.05 | 0.07 | -0.03 | 0.00 |
| PC34 vegetation | 0.01 | -0.01 | 0.11 | 0.00 | -0.12 | -0.08 | 0.14 | 0.05 | 0.12 | -0.05 | -0.15 | 0.19 | -0.23 | -0.31 | -0.02 | 0.11 | 0.65 | 0.15 | 0.18 | -0.02 | -0.11 | -0.04 | -0.11 | 0.04 | -0.04 | 0.08 | -0.01 | -0.13 | -0.02 | -0.04 | -0.02 | 0.05 | 0.05 | -0.18 | -0.05 | 0.04 | -0.01 | 0.18 | -0.15 | 0.00 | 0.01 | 0.00 | 0.10 | 0.04 | 0.24 | 0.09 | 0.02 | 0.00 | 0.02 | -0.01 | -0.01 | 0.04 |
| PC35 vegetation | 0.01 | -0.04 | -0.01 | -0.12 | 0.12 | -0.10 | 0.01 | 0.00 | 0.08 | 0.10 | -0.16 | -0.08 | 0.14 | 0.08 | 0.21 | 0.12 | 0.17 | 0.15 | 0.07 | -0.20 | -0.23 | 0.06 | 0.35 | 0.06 | 0.18 | -0.37 | -0.10 | 0.24 | -0.09 | -0.17 | 0.00 | 0.05 | 0.29 | -0.07 | 0.05 | 0.15 | 0.11 | -0.22 | 0.20 | 0.09 | 0.08 | -0.10 | 0.04 | 0.12 | -0.03 | -0.03 | 0.04 | 0.08 | 0.10 | 0.04 | -0.01 | -0.02 |
| PC36 vegetation | 0.00 | 0.00 | 0.09 | 0.06 | -0.13 | 0.04 | 0.06 | 0.07 | 0.03 | 0.05 | 0.12 | 0.03 | 0.14 | 0.11 | 0.16 | 0.04 | -0.02 | 0.01 | 0.10 | -0.02 | -0.01 | 0.11 | 0.48 | 0.15 | -0.16 | 0.36 | -0.28 | -0.12 | 0.09 | 0.21 | -0.20 | -0.35 | 0.22 | -0.06 | -0.06 | -0.06 | -0.14 | -0.06 | 0.00 | -0.14 | -0.09 | 0.09 | 0.00 | -0.05 | 0.16 | 0.08 | -0.03 | -0.04 | -0.01 | 0.00 | 0.00 | 0.00 |
| PC37 vegetation | 0.01 | 0.01 | 0.07 | -0.09 | 0.04 | 0.02 | -0.13 | 0.00 | -0.02 | 0.02 | 0.08 | 0.10 | 0.18 | 0.25 | 0.08 | -0.12 | 0.26 | 0.10 | 0.16 | -0.06 | -0.09 | -0.21 | -0.18 | -0.02 | 0.02 | -0.35 | -0.06 | -0.08 | 0.00 | 0.09 | -0.03 | -0.43 | -0.30 | 0.21 | -0.05 | -0.01 | -0.24 | -0.17 | -0.24 | 0.00 | -0.02 | 0.06 | -0.10 | -0.05 | -0.03 | 0.02 | -0.02 | 0.10 | 0.04 | 0.01 | 0.00 | 0.00 |
| limestone | 0.00 | 0.06 | -0.19 | 0.12 | 0.22 | -0.52 | -0.08 | -0.25 | 0.07 | 0.13 | -0.11 | 0.02 | 0.01 | -0.02 | 0.02 | 0.01 | -0.01 | 0.00 | 0.00 | 0.01 | 0.00 | 0.01 | 0.00 | 0.00 | 0.00 | 0.00 | 0.00 | 0.00 | 0.00 | 0.00 | 0.00 | -0.01 | 0.00 | 0.00 | 0.00 | 0.00 | -0.02 | 0.00 | -0.02 | -0.08 | -0.04 | 0.58 | 0.28 | -0.31 | -0.01 | -0.04 | 0.03 | -0.02 | -0.06 | -0.03 | 0.00 | -0.02 |
| serpentinite | 0.02 | -0.01 | 0.03 | 0.00 | 0.04 | 0.02 | 0.03 | -0.03 | 0.07 | 0.06 | -0.04 | -0.28 | 0.38 | 0.38 | 0.10 | -0.06 | 0.13 | 0.02 | 0.03 | -0.03 | 0.00 | 0.00 | -0.04 | 0.00 | -0.01 | 0.01 | 0.01 | -0.04 | 0.00 | -0.03 | 0.03 | 0.07 | 0.02 | -0.02 | 0.02 | -0.01 | 0.12 | 0.75 | -0.04 | -0.06 | 0.07 | -0.01 | 0.02 | 0.00 | 0.00 | 0.00 | 0.01 | 0.00 | 0.01 | 0.00 | 0.00 | 0.00 |
| is Hokkaido | -0.06 | -0.57 | 0.02 | -0.02 | 0.12 | -0.03 | 0.00 | 0.06 | 0.03 | 0.14 | 0.10 | -0.02 | -0.02 | 0.00 | -0.01 | -0.01 | 0.00 | 0.00 | 0.00 | 0.00 | 0.00 | 0.00 | 0.00 | 0.00 | 0.00 | 0.00 | 0.00 | 0.00 | 0.00 | 0.00 | 0.00 | 0.00 | 0.00 | 0.01 | 0.00 | 0.00 | 0.01 | -0.01 | 0.00 | 0.02 | -0.02 | 0.02 | -0.01 | 0.02 | -0.02 | 0.05 | 0.02 | 0.01 | -0.15 | -0.43 | 0.21 | 0.60 |
| is oceanic island | -0.04 | 0.04 | -0.29 | 0.09 | 0.46 | 0.31 | -0.20 | -0.18 | -0.13 | -0.03 | 0.05 | 0.00 | 0.00 | 0.00 | 0.00 | 0.00 | -0.01 | 0.00 | 0.00 | 0.00 | 0.00 | 0.00 | 0.00 | 0.00 | 0.00 | -0.01 | 0.00 | 0.00 | 0.00 | 0.00 | 0.00 | 0.00 | -0.01 | 0.01 | 0.00 | 0.01 | -0.01 | -0.01 | -0.01 | 0.01 | 0.04 | -0.01 | 0.06 | 0.19 | 0.67 | 0.05 | 0.07 | -0.02 | -0.03 | -0.02 | 0.01 | -0.02 |

### Data distibution against the environmental variables used

#### Climatic variables

```
> p_climate = (
+   (p7+p8+p9 +plot_layout(guides = "collect", ncol = 2))
+   & labs(x = NULL, y = NULL)
+   & theme(plot.title = element_text(hjust = 1, vjust = -8))
+   )
> grid.arrange(
+   patchworkGrob(p_climate), 
+   top = NULL,
+   left = ylab, 
+   bottom = str_c(xlab,"\nS1-1 Fig. all climate variables used in SDMs")
+   )
```

#### Land use variables

```
> p_landuse = (
+   (p1+p2+p3+p4+p5+p6+ 
+      plot_layout(guides = "collect", ncol = 3)
+    )
+   & labs(x = NULL, y = NULL)
+   & theme(plot.title = element_text(hjust = 1, vjust = -8))
+   )
> grid.arrange(
+   patchworkGrob(p_landuse), 
+   left = ylab,
+   bottom = str_c(xlab, "\nS1-2 Fig. all land use variables used in SDMs"),
+   top = NULL
+   )
```

#### Geological variables

Note that longitude and latitude were not used in the SDMs.

```
> p_geo = (
+   ((p10/p11)|(p51/p52)|(p53/p54)) #p50
+   & labs(x = NULL, y = NULL)
+   & theme(plot.title = element_text(hjust = 1, vjust = -8))
+   )
> grid.arrange(
+   patchworkGrob(p_geo), 
+   left = ylab,
+   bottom = str_c(xlab, "\nS1-3 Fig. all geological variables used in SDMs"),
+   top = NULL
+   )
```

#### Vegetation variables

```
> p_geo = (
+   (p12+p13+p14+p15+p16+p17+p18+p19+p20+p21+
+   p22+p23+p24+p25+p26+p27+p28+p29+p30+p31+
+   p32+p33+p34+p35+p36+p37+p38+p39+p40+p41+
+   p42+p43+p44+p45+p46+p47+p48+
+     plot_layout(guides = "collect", ncol = 3))
+   & labs(x = NULL, y = NULL)
+   & theme(plot.title = element_text(hjust = 1, vjust = -8))
+   )
> grid.arrange(
+   patchworkGrob(p_geo), 
+   left = ylab,
+   bottom = str_c(xlab, "\nS1-4 Fig. all geological variables used in SDMs"),
+   top = NULL
+   )
```
