## Supplementary File 2 for "Boosting biodiversity monitoring using smartphone-driven, rapidly accumulating community-sourced data"

### List of GBIF occurrence doi

*GBIF occurrences were downloaded at each order on 06 July 2023: DOI10.15468/dl.2av9qf ; DOI10.15468/dl.2cmsas ; DOI10.15468/dl.2huakz ; DOI10.15468/dl.2kbfw4 ; DOI10.15468/dl.2zuxyj ; DOI10.15468/dl.32sraw ; DOI10.15468/dl.34nv8h ; DOI10.15468/dl.39v53g ; DOI10.15468/dl.3b77pn ; DOI10.15468/dl.3hwgta ; DOI10.15468/dl.3ss8vz ; DOI10.15468/dl.3z8284 ; DOI10.15468/dl.44r7sv ; DOI10.15468/dl.49g4h6 ; DOI10.15468/dl.4a4u9f ; DOI10.15468/dl.4fc29k ; DOI10.15468/dl.4tdngj ; DOI10.15468/dl.4tq9hk ; DOI10.15468/dl.4vz9a4 ; DOI10.15468/dl.4xnud6 ; DOI10.15468/dl.4yk74w ; DOI10.15468/dl.4zwq9c ; DOI10.15468/dl.566ebu ; DOI10.15468/dl.5b6ngk ; DOI10.15468/dl.5cjvcx ; DOI10.15468/dl.5ktfys ; DOI10.15468/dl.5vsqq9 ; DOI10.15468/dl.5vzzxw ; DOI10.15468/dl.5zrynk ; DOI10.15468/dl.6at6x4 ; DOI10.15468/dl.6avq4x ; DOI10.15468/dl.6cgxtc ; DOI10.15468/dl.75rscg ; DOI10.15468/dl.76vwv2 ; DOI10.15468/dl.7s5gmg ; DOI10.15468/dl.7tjddr ; DOI10.15468/dl.7ubqp8 ; DOI10.15468/dl.7wq6td ; DOI10.15468/dl.7yrxgv ; DOI10.15468/dl.7z4vz2 ; DOI10.15468/dl.7zk475 ; DOI10.15468/dl.855fps ; DOI10.15468/dl.892guv ; DOI10.15468/dl.8setn5 ; DOI10.15468/dl.8th3fd ; DOI10.15468/dl.8wvr75 ; DOI10.15468/dl.93wd8q ; DOI10.15468/dl.93z9zc ; DOI10.15468/dl.96fb9d ; DOI10.15468/dl.9axnge ; DOI10.15468/dl.9bqd88 ; DOI10.15468/dl.9e7e7d ; DOI10.15468/dl.9jcj3d ; DOI10.15468/dl.9m42rm ; DOI10.15468/dl.9mdk8n ; DOI10.15468/dl.9p7p54 ; DOI10.15468/dl.9uemr3 ; DOI10.15468/dl.9v7hxv ; DOI10.15468/dl.9va7q5 ; DOI10.15468/dl.9wu8ry ; DOI10.15468/dl.9xtdyu ; DOI10.15468/dl.a2fr7h ; DOI10.15468/dl.a5zh92 ; DOI10.15468/dl.a8694f ; DOI10.15468/dl.aedkqc ; DOI10.15468/dl.ahdvg5 ; DOI10.15468/dl.b4je8b ; DOI10.15468/dl.bb7k2w ; DOI10.15468/dl.bkbj9r ; DOI10.15468/dl.bp3ndm ; DOI10.15468/dl.brbrr3 ; DOI10.15468/dl.bt2ypa ; DOI10.15468/dl.bw6h94 ; DOI10.15468/dl.bzb483 ; DOI10.15468/dl.cgc8yt ; DOI10.15468/dl.cj6ujy ; DOI10.15468/dl.ckeuvm ; DOI10.15468/dl.cm4x2b ; DOI10.15468/dl.cn2zqn ; DOI10.15468/dl.cnpybg ; DOI10.15468/dl.cr6jy3 ; DOI10.15468/dl.cs9cvq ; DOI10.15468/dl.ct8hg6 ; DOI10.15468/dl.czkyce ; DOI10.15468/dl.d3wb86 ; DOI10.15468/dl.db8uex ; DOI10.15468/dl.ddh496 ; DOI10.15468/dl.de9mq5 ; DOI10.15468/dl.dewykn ; DOI10.15468/dl.dfvtr3 ; DOI10.15468/dl.dqecub ; DOI10.15468/dl.dve7qt ; DOI10.15468/dl.dy8euw ; DOI10.15468/dl.e76gdv ; DOI10.15468/dl.ebah9w ; DOI10.15468/dl.er8sd2 ; DOI10.15468/dl.esnvqy ; DOI10.15468/dl.f2u2fk ; DOI10.15468/dl.f4jhnc ; DOI10.15468/dl.f6r2uk ; DOI10.15468/dl.f759y9 ; DOI10.15468/dl.f9p655 ; DOI10.15468/dl.fexq22 ; DOI10.15468/dl.fgrq2f ; DOI10.15468/dl.frm4vg ; DOI10.15468/dl.fuvg9w ; DOI10.15468/dl.g9q8r7 ; DOI10.15468/dl.gaa6gm ; DOI10.15468/dl.gcnmgx ; DOI10.15468/dl.gdrv3r ; DOI10.15468/dl.ggdjvv ; DOI10.15468/dl.gj7u3p ; DOI10.15468/dl.gtxra2 ; DOI10.15468/dl.h33ube ; DOI10.15468/dl.h79vnk ; DOI10.15468/dl.h96p3j ; DOI10.15468/dl.hj8kyh ; DOI10.15468/dl.hjjezh ; DOI10.15468/dl.hkaf2e ; DOI10.15468/dl.hpmhfg ; DOI10.15468/dl.hrtsyq ; DOI10.15468/dl.hygmvs ; DOI10.15468/dl.je3fvj ; DOI10.15468/dl.jh95fq ; DOI10.15468/dl.jhpgp9 ; DOI10.15468/dl.jnup28 ; DOI10.15468/dl.juj8cn ; DOI10.15468/dl.jw6yme ; DOI10.15468/dl.kew5f9 ; DOI10.15468/dl.kff3wf ; DOI10.15468/dl.m9twve ; DOI10.15468/dl.mfrq4p ; DOI10.15468/dl.mtbaxw ; DOI10.15468/dl.mzvkf7 ; DOI10.15468/dl.na89yg ; DOI10.15468/dl.ndyscc ; DOI10.15468/dl.nq62ve ; DOI10.15468/dl.nutb8c ; DOI10.15468/dl.nxqy2a ; DOI10.15468/dl.p5gw4t ; DOI10.15468/dl.p5szjt ; DOI10.15468/dl.p7rxga ; DOI10.15468/dl.pazc53 ; DOI10.15468/dl.pbhk5b ; DOI10.15468/dl.pfnv5e ; DOI10.15468/dl.pjx7bg ; DOI10.15468/dl.pvfbnq ; DOI10.15468/dl.pwyz6j ; DOI10.15468/dl.q2br6n ; DOI10.15468/dl.q4gunv ; DOI10.15468/dl.q4j76s ; DOI10.15468/dl.q73x7a ; DOI10.15468/dl.q9y2x5 ; DOI10.15468/dl.qbhyan ; DOI10.15468/dl.qqzfkq ; DOI10.15468/dl.qvyk6k ; DOI10.15468/dl.qyswun ; DOI10.15468/dl.rjsuxy ; DOI10.15468/dl.s2uqms ; DOI10.15468/dl.sdvshe ; DOI10.15468/dl.sdz9c2 ; DOI10.15468/dl.sgc9vz ; DOI10.15468/dl.sh7etj ; DOI10.15468/dl.smhu6k ; DOI10.15468/dl.syn437 ; DOI10.15468/dl.t2y3bn ; DOI10.15468/dl.tm335t ; DOI10.15468/dl.tnehdb ; DOI10.15468/dl.trpwse ; DOI10.15468/dl.txbnha ; DOI10.15468/dl.ty8wua ; DOI10.15468/dl.tzsrxq ; DOI10.15468/dl.u8z38k ; DOI10.15468/dl.u9uazh ; DOI10.15468/dl.udnxp7 ; DOI10.15468/dl.uegp3a ; DOI10.15468/dl.uj5za2 ; DOI10.15468/dl.uzd7pz ; DOI10.15468/dl.vsrmm4 ; DOI10.15468/dl.vwmgn3 ; DOI10.15468/dl.w2txce ; DOI10.15468/dl.w4upfs ; DOI10.15468/dl.w586u3 ; DOI10.15468/dl.wcgtbb ; DOI10.15468/dl.wcqf67 ; DOI10.15468/dl.wms5aj ; DOI10.15468/dl.wnfcjg ; DOI10.15468/dl.wpnsb2 ; DOI10.15468/dl.ws6s5j ; DOI10.15468/dl.wtkjbq ; DOI10.15468/dl.wxpqyz ; DOI10.15468/dl.x95raj ; DOI10.15468/dl.xbqc6y ; DOI10.15468/dl.xermz5 ; DOI10.15468/dl.xkqch8 ; DOI10.15468/dl.xrmt5h ; DOI10.15468/dl.y5knh6 ; DOI10.15468/dl.y5qjam ; DOI10.15468/dl.yamk7y ; DOI10.15468/dl.yey48w ; DOI10.15468/dl.yhfx9y ; DOI10.15468/dl.yqm2uc ; DOI10.15468/dl.yrw7nm ; DOI10.15468/dl.yzxa75 ; DOI10.15468/dl.z68efe ; DOI10.15468/dl.z73afv ; DOI10.15468/dl.z7emth ; DOI10.15468/dl.z8y7fy ; DOI10.15468/dl.zgyeak ; DOI10.15468/dl.zmjrg6 ; DOI10.15468/dl.zpwtjt ; DOI10.15468/dl.ztdqas ; DOI10.15468/dl.zuudhh ; DOI10.15468/dl.zv5yhq

### List of literatures

Misao Tatewaki. A List of Plants col1eted in the Wakayama University Experimental Fosest. (1). Research bulletins of the College Experiment Forests, College of Agriculture, Hokkaido Imperial University. 1932;7: 131–180.

Taya I, Sokabe Y, Harada A, Yoshida H, Mikazuki A, Tawa K, et al. A report of the biological research on butterflies, amphibians, reptiles and wild birds in the Yaeyama Islands of Okinawa prefecture, Japan. Memoirs of the Faculty of Agriculture of Kinki University. 2013; 299–307.

Tsuji Y, Kondo H. Ant (Hymenoptera: Formicidae) fauna of Noichi zoological park of kochi prefecture, Japan. Bulletin of the Shikoku Institute of Natural History. 2022;15: 46–52. doi:[10.32250/sinh.15.0_46](https://doi.org/10.32250/sinh.15.0_46)

Yasuhiko Konno, Yasuyuki Watanabe, Nobuki Yasuda. Aquatic Insect Fauna of the High Altitude Zone in Daisetsu Mountains. Bulletin of Sounkyo Visitor Center. 2001;21: 33–39.

Oba Shinya, Iguchi Keiichirou. Aquatic insects found in fallow fields in Tatsugo-cho, Oshima-gun in the spring of 2020. Nature of Kagoshima. 2021; 397–398.

Kaoru Haga. Beetles collected from Shiretoko Pass. Sylvicola. 1996; 21–31.

Takanori Sato, Shigehiro Nakabayashi, Nobuyuki Narumi, Yukihiro Kohmatsu. Breeding conditions of the Siberian salamander Salamaηdrella keyserlingii in Kushiro Mire. Memoirs of the Kushiro City Museum. 1999;23: 25–31. doi:[10.24484/sitereports.118607-67309](https://doi.org/10.24484/sitereports.118607-67309)

Matsumoto T, Sato S, Inoue T. Butterfly fauna of Shikoku Research Center, Forestry and Forest Products Research Institute. Bulletin of the Forestry and Forest Products Research Institute. 2013;12: 111–124.

Masahiko Nakatani. Coleoptera fauna in an urban area of Kushiro, Hokkaido. Sylvicola. 2010; 83–86.

Masahiko Nakatani, Ken-ichi Matsumoto. Coleoptera fauna of Kaminoko-ike Pond. Kiyosato-cho, Eastern Hokkaido. Sylvicola. 2009; 11–16.

Masahiro Mitsuo. Comparison of the plant associations in Shiroyama, Kagoshima city between 1974 and 1991. Bulletin of the Kagoshima Prefectural Museum. 1992; 43–49.

Takumi Komatsu. Distributional records of Scarabaeoidea (Coleoptera) from Kakeroma-jima Island, Amami Islands, central Ryukyus, southwestern Japan. Fauna Ryukyuana. 63: 21–28.

Shuichi Ikudome. Ecological Studies on the Wild Bee・ Fauna in the Rural Area on Yaku-shima, Kagoshima Prefecture, Nippon (Hymenoptera, Apoidea). Bulletin of Kagoshima Women’s Junior College Kagoshima. 2005;40: 1–20.

Yoko Nishikawa, Masami Miyaki, Shigehisa Hori. Effects of Pinus pumila on the distribution of alpine plants and Callianthemum miyabeanum at Mt. Apoi. Research Report of Hokkaido Institute of Enviromental Sciences. 1993; 89–95.

Tomiyama H. Ferns of Kumejima. Kumejima Comprehensive Survey Report Nature, History, Folklore, Koko, Arts and Crafts, Architecture, Okinawa Prefectural Museum. 1994; 12–25.

Shinichi Niwa, Osamu Watanabe, Noriyuki Watanabe. Flora of Tanetomi Wetland, Rishiri Island, Hokkaido. Rishiri Studies. 2001; 69–74.

Nobuki Yasuda, Masahiko Sato. Insect Faunal Survey of Is. Rishiri and Is. Rebun and Sarobetsu Field - The Vertical Distribution of Ground Beetle Communities in Mt. Rebun, Is. Rebun, Hokkaido. Rishiri Studies. 1992; 11–25.

Nobuki Yasuda, Eiji Nishitani, Masahiko Sato. Insect Faunal Survey of Is. Risiri and Is. Rebun - The Vertical Distribution of Ground Beetle Communities in Mt. Risiri, Is. Risiri, Hokkaido. Rishiri Studies. 1991; 13–28.

Yamazaki J, Matsumura M, Kohama T, Osada M, Nobayashi S. Insect list of Iheyajima Island and Nohojima Island. Reprinted from Survey Reports on Natural History, History and Culture of Izenajima and Iheyajima Islands, Okinawa Prefectural Museum and Art Museum. 2019; 25–36.

Moriyama T, Kanai K. Insects collected on Kuchino-shima,Nakano-shima and Swanose-jima (Tokara Islands) in 2015. Bulletin of the Kagoshima Prefectural Museum. 2016; 57–66.

Kenichi Kanai. Insects collected on Kuchinoerabu-jima (Osumi Islands) in 2016. Bulletin of the Kagoshima Prefectural Museum. 2017; 15−23.

Ken-ichi Matsumoto, Masahiko Nakatani. Insects fauna of Akan National Park IV. Hokkaido. Sylvicola. 2010; 11–20.

Nemu Tsuji, Norihito Muramatsu, Masahiko Nakatani, Isamu Nakamura, Tadaaki Sato. Insects of Okukussharo Onsen in Tsubetsu Town, Hokkaido. Sylvicola. 2010; 1–10.

Nakamine K. Insects of Taira-jima and Nakano-shima, in Tokara-retto, Kagoshima Prefecture, surveyed in autumn, 2007. Bulletin of the Kagoshima Prefectural Museum. 2008; 83–92.

Koji Nakamine. Insects recorded in 2005 in Koshiki Islands, Kagoshima Prefecture (2). Bulletin of the Kagoshima Prefectural Museum. 2007; 79–87.

Koji Nakamine. Insects recorded in 2005 in Koshikijima Islands,Kagoshima Prefecture(1). Bulletin of the Kagoshima Prefectural Museum. 2006; 38–55.

Koji Nakamine. Insects recorded in August,2005 in Take-shima,Mishima, Kagoshima Prefecture. Bulletin of the Kagoshima Prefectural Museum. 2006; 56–62.

Koji Nakamine. Insects recorded in September, 2004 in Uke-j ima,Amami-islands, Kagoshima Prefecture. Bulletin of the Kagoshima Prefectural Museum. 2006; 63–67.

Kunikichi Yoshida. Moth Fauna of Tomakomai Experiment Forest of Hokkaido University. Research Bulletin of the Hokkaido University Forests. 1976;33: 457–480.

Naomi Miyazaki. Plant species diversity in Man-made forest “Obihiro no Mori” and its interaction with local dwellers. Master’s Course of Ecological Science of Obihiro University. 2011.

Kido N. Plants of Kodakarajima. Quarterly journal of welfare society. 2014;33: 72–87.

Shizuo Sako. Preliminary Reports on the Flowering Plants and Ferns hither-to known in the Takakuma Experimental Forest of Kagoshima University. Res Bull Kagoshima Univ For. 1968;1: 38–125.

Satoshi Ohya. Record of plant collection on Tokuno-shima Island, Kagoshima Prefecture. Bulletin of the Kagoshima Prefectural Museum. 2008; 59–64.

Shigeko Fukumoto, Masaki Yamane. Records of ants from Kikai-Jima, the Amami Islands, Kagoshima Prefecture, Japan (Hymenoptera, Formicidae). Nature of Kagoshima. 2020; 509–514.

Daiki Uchida, Tadashi Kitano, Shingo Sano, Ryouichi Furuhata, Kohei Watanabe. Records of aquatic Hemiptera from Kakeroma-jima Island, Amami Islands, Japan. Rostria : Transactions of the Hemipterological Society of Japan. 2021; 30–34.

Kawanishi Motohiro, Satou Haruhi, Aiba Shin-ichiro, Fujita Shiho, Ugawa Shin, Eimura Naoko, et al. Relationships between clear cutting history and/or microtopography and species diversity in the laurel forest of Amami-Oshima Island, a World Natural Heritage candidate, Kagoshima, Japan. Annual Report of Pro Natura Foundation Japan. 2021;30: 6–24. doi:[10.32215/pronatura.30.0_6](https://doi.org/10.32215/pronatura.30.0_6)

Hisamitsu A, Sokabe Y, Torii N, Kuwabara T, Suzuki K, Yamamoto Y, et al. Report of biological research on Lepidoptera in Yaeyama Islands, Oldnawa prefecture, Japan. Memoirs of the Faculty of Agriculture of Kinki University. 2011; 139–149.

Sakuma T, Yamashiro S, Moriguchi Y, Ishikawa S, Miyake N. Species diversity of the plant communities and flora of Naruyama area, Ino town, Kochi Prefecture. Bulletin of the Shikoku Institute of Natural History. 2006;3: 78–85. doi:<https://doi.org/10.32250/sinh.3.0_78>

Tamotsu Hattori, Daisuke Tochimoto, Koji Iwakiri, Noriko Minamiyama, Yoshinobu Hashimoto. Species richness and species composition of epiphytes in a lucidophyllous forest on Mt. Kurino-dake, Kagoshima Prefecture. Humans and Nature. 2007;18: 29–38. doi:[10.24713/hitotoshizen.18.0_29](https://doi.org/10.24713/hitotoshizen.18.0_29)

Tamotsu Hattori, Daisuke Tochimoto, Noriko Minamiyama, Yoshinobu Hashimoto, Yoshihiro Sawada, Hiroaki Ishida. Species richness and species composition of vascular epiphytes in the lucidophyllous forests in southern Kyushu. Vegetation Science. 2009;26: 49–61. doi:[10.15031/vegsci.26.49](https://doi.org/10.15031/vegsci.26.49)

Masatoshi Takakuwa, FUjita Hiroshi. Spring Research on Insect Fauna near Hananoego of the Montane Zone of Yaku-shima Island, Southwest Japan. Bulletin of the Kanagawa Prefectrural Museum Natural science. 2010; 35–38.

Hattori T, Iwakiri K, Minamiyama N, Kuroki S, Kuroda A. Structure, species richness, and species composition of the shrine forest in Miyazaki-Jingu, Miyazaki Prefecture. Japanese Journal of Conservation Ecology. 2010;15: 47–59. doi:<https://doi.org/10.18960/hozen.15.1_47>

Jinshi Terada, Motohiro Kawanishi, Koshiro Kubo. The Dune Vegetation of Honmura Beach in Tanegashima Island. Bulletin of the Kagoshima Prefectural Museum. 2014; 1–26.

Norio Murano. The Flora and Fauna of Nopporo Forest Park. Journal of Rakuno Gakuen University Natural science. 2000;25: 133–200.

Kenji Hatada. The insects fauna of Satsuma-Io-jima Isls., Kagoshima Pref. Bulletin of the Kagoshima Prefectural Museum. 1990; 9–13.

Ie Village Board of Education. The Picture Guide Book of Plants in Ie Island. 2002.

Hidetaka Umata, Masami Matsumoto. The plants of Takakuma Experimental Forest －Supplement (2)－. Res Bull Kagoshima Univ For. 2006;34: 81–82.

Hidetaka Umata, Munetaka Nishi, Katsuo Maruyama. The plants of Takakuma Experimental Forest －Supplement (3)－. Res Bull Kagoshima Univ For. 2007;35: 61–63.

Morita Y. The Report of the Plant Collection on Kuchino-shima in the Takara Islands,Kagoshima Prefecture. Bulletin of the Kagoshima Prefectural Museum. 2004; 55–60.

Yasuo Morita, Katahira Takao. The report of the plant collection on Shimokoshiki-jima Island, Kagoshima Prefecture. Bulletin of the Kagoshima Prefectural Museum. 2006; 12–21.

Morita Y. The report of the plant collection on Suwanosezima in the Takara Islands, Kagoshima Prefecture. Bulletin of the Kagoshima Prefectural Museum. 2001; 25–31.

Yasuo Morita. The Report of the Plant Collection on Tanega-shima, Kagoshima Prefecture. Bulletin of the Kagoshima Prefectural Museum. 2004; 61–73.

Terada J, Kawanishi M, Yamazaki J. The Vegetation of Kubayama covered Livistona chinensis forest Iheyajima Island, Okinawa Prefecture. Reprinted from Survey Reports on Natural History, History and Culture of Izenajima and Iheyajima Islands, Okinawa Prefectural Museum and Art Museum. 2019; 77–94.

Niiro Y, Shinjo K. Vegetation of lheya-Izena Shoto. Diss University of Ryukyus. 1959; 81–105.

Shikoku Institute of Natural History. Yokonami Peninsula Organism Comprehensive Survey Report. Shikoku Institute of Natural History; 2018. Available:<http://www.lutra.jp/>
